# fMRI repetition suppression reveals no sensitivity to trait judgments from faces along the ventral visual stream or in the theory-of-mind network

**DOI:** 10.1101/351783

**Authors:** Emily E. Butler, Rob Ward, Paul E. Downing, Richard Ramsey

## Abstract

The human face cues a wealth of social information, but the neural mechanisms that underpin social attributions from faces are not well known. In the current fMRI experiment, we used repetition suppression to test the hypothesis that populations of neurons in face perception and theory-of-mind neural networks would show sensitivity to faces that cue distinct trait judgments. Although faces were accurately discriminated based on associated traits, our results showed no evidence that face or theory-of-mind networks showed repetition suppression for face traits. Thus, we do not provide evidence for population coding models of face perception that include sensitivity to high and low trait features. Due to aspects of the experimental design, which bolstered statistical power and sensitivity, we have reasonable confidence that we could detect effects of a moderate size, should they exist. The null findings reported here, therefore, add value to models of neural organisation in social perception by showing instances where effects are absent or small. To test the generalisability of our findings, future work should test different types of trait judgment and different types of facial stimuli, in order to further probe the neurobiological bases of impression formation based on facial appearance.

## Introduction

Faces signal information that guide social interactions (Emery, 2000). Although complex social signals such as emotional states, trait characteristics, and attentional focus are readily perceived from faces (Jack & Schyns, 2017; Todorov et al., 2015), the neural mechanisms that process social dimensions of face perception remain unclear. Here, in a functional magnetic resonance imaging (fMRI) experiment, we use repetition suppression to investigate the neural representation of how trait inferences are arrived at during social perception.

The majority of neuroscience research on face perception has focused on detection and recognition of identity and emotion. This research has identified face-selective patches of cortex that respond more to viewing faces than other categories of objects such as houses and cars (Duchaine & Yovel, 2015; Haxby et al., 2000; Kanwisher et al., 1997). Key regions in the face perception network include the fusiform face area (FFA; Kanwisher et al., 1997), occipital face area (OFA; Gauthier et al., 2000) and posterior superior temporal sulcus (pSTS; Allison et al., 2000; Pitcher et al., 2011). These three nodes along the ventral visual stream are suggested to perform core visual analyses of facial features, but also interact with extended circuits in anterior cortex for more elaborate representations of identity and emotional valence (Duchaine & Yovel, 2015; Haxby et al., 2000; Kanwisher, 2010).

Face recognition is important for initiating social interactions, but faces cue much more than the mere presence of a social agent. Indeed, impressions of others are partly formed on the basis of stable, non-emotional aspects of facial appearance (Todorov et al., 2015; Zebrowitz, 2011). As such, there is interplay between the perception of facial features and the formation of character impressions (Jack & Schyns, 2017). Models of social impressions from faces have been developed that include dimensions of valence/trustworthiness, dominance and attractiveness (Todorov et al., 2008; Sutherland et al., 2013; Wang et al., 2016). However, there is currently little known regarding the neural bases of such impression formation. For example, faces that cue social evaluations of trustworthiness and attractiveness have been associated with responses in the amygdala and ventral striatum, which have been thought to index the reward value and typicality of faces (Bzdok et al., 2011; Mende-Siedlecki et al., 2013; Said et al., 2010; 2011; Todorov et al., 2013). Additionally, behavioural research has shown that personality characteristics such as extraversion are accurately perceived from static facial features (Borkenau & Liebler, 1992; Borkenau et al., 2009; Kramer & Ward, 2010; Penton-Voak et al., 2006). However, beyond brain circuits associated with reward, little is currently known regarding the neural architecture supporting personality judgments that are cued during face perception.

Research investigating trait judgments has primarily focused on reading statements, rather than faces (Uleman et al., 2008). For example, reading trait-diagnostic statements, such as “she gave money to charity”, engages the theory-of-mind (ToM) network more than trait-neutral statements such as “she sharpened her pencil” (Heleven et al., 2017; Heleven & Van Overwalle, 2016; Ma et al., 2014; Mitchell et al., 2002; 2005; Van Overwalle et al., 2016). The ToM network is engaged when attributing mental states such as beliefs, desires and attitudes to others, as well as judging character and is thought to be central to understanding social cognition (van Overwalle, 2009; Frith & Frith, 1999). The ToM network is largely distinct from the face perception network with key nodes covering temporoparietal junction, medial prefrontal cortex, temporal poles and precuneus (van Overwalle, 2009; Frith & Frith, 1999; Saxe & Kanwisher, 2003). However, the potential role of the ToM network in forming impressions based on facial appearance has not been studied in depth. As such, the cognitive and neural systems that identify perceptual features and link them to trait judgments are not well known (Over & Cook, 2018). The current study, therefore, investigates the hypothesis that impression formation from faces relies on a distributed neural architecture that spans the face perception and ToM neural networks.

In the current fMRI study, we addressed the extent to which face perception and ToM networks contribute to forming impressions based on facial appearance. The experiment used a repetition suppression (RS) design (Grill-Spector, Henson, & Martin, 2006; Barron et al., 2016). RS designs measure a reduced BOLD response following a repeated stimulus feature and a release from suppression following a novel stimulus feature. Compared to conventional subtraction designs, which can show if a brain region shows magnitude differences between conditions, RS studies hold the potential to study neural processes at the level of neural populations within a given brain region. A brain region that shows RS, therefore, can allow inferences about the organisation of underlying neural populations (Barron et al., 2016; Figure 1). We created face stimuli that cued high and low trait judgments and showed these stimuli to participants in a sequence that created novel and repeated events. To identify functional regions of interest, we used established face and ToM localiser tasks and to bolster statistical power we used an analysis pipeline that has been demonstrated to exhibit high functional resolution and sensitivity (Julian et al., 2012; Nieto-Castanon & Fedorenko, 2012). If the face and ToM networks are engaged in forming impressions based on facial features in the manner that we predict, we would expect to observe repetition suppression for face traits in both networks.

**Figure 1.**
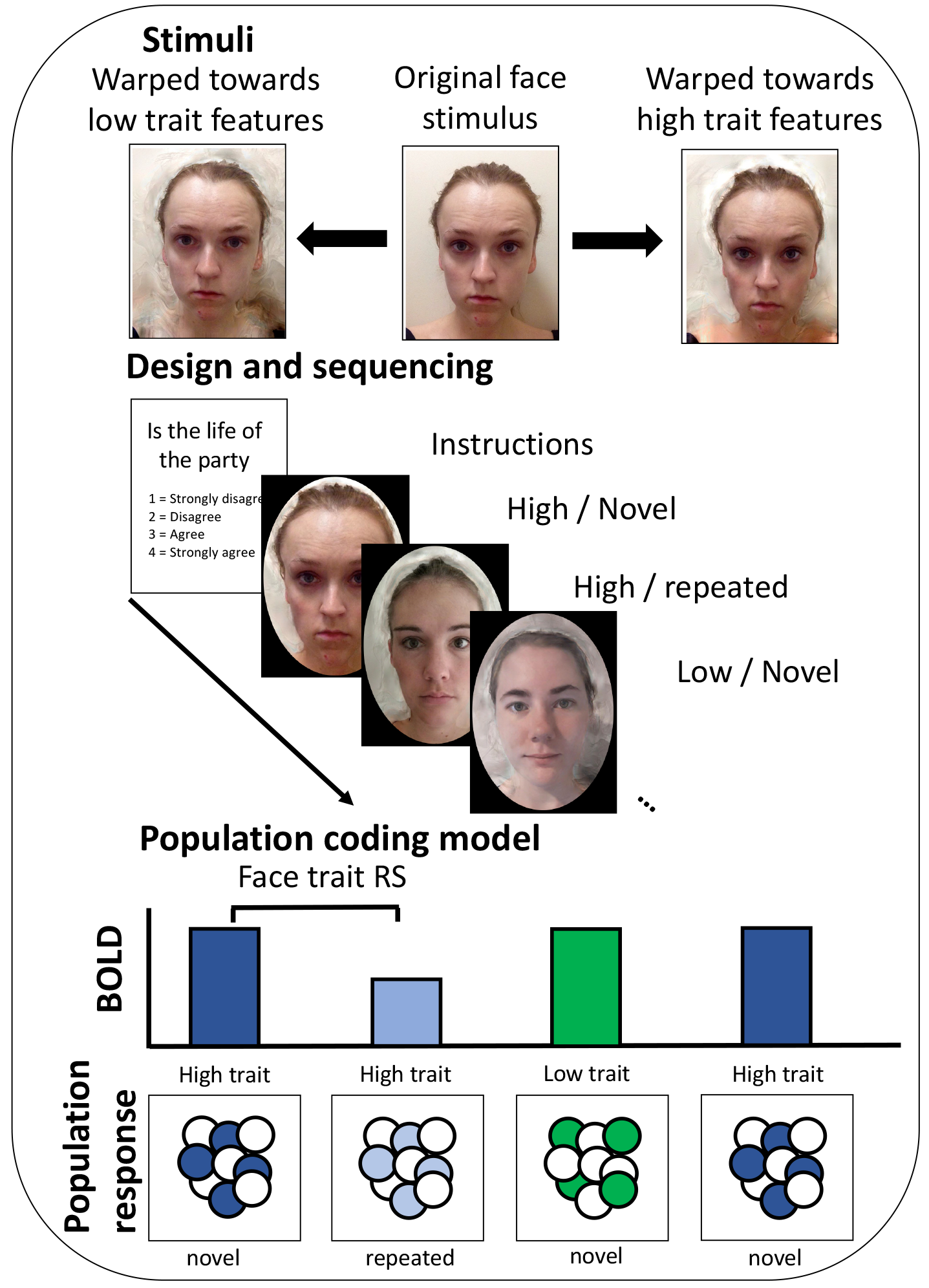
Method and design. A) Individual face images were transformed towards high and low composite templates of trait variables (Extraversion, Agreeableness, Neuroticism, Physical health). The example shown is extraversion. The images used are for illustrative purposes and were not used in the experiment. B) During scanning, each block began with an instruction screen, which provided a statement and a reminder of the ratings scale. On each subsequent trial, participants had to make a judgment based on the face presented. As such, all trials in a mini-block were from the same category (e.g., extraversion), but all trials showed a different individual. C) An illustration of the population coding model of face perception that the repetition suppression design was testing. High and low trait features are presented in blue and green, respectively. Novel and repeated trials are presented in darker and lighter colours, respectively.

## Method

### Participants

Twenty-eight participants completed the experiment (14 female; M_age_=23.96, SD=5.52). All participants received a monetary reimbursement (£15), had normal or corrected-to-normal vision and gave informed consent according to the local ethics guidelines.

### Stimuli and experimental tasks

#### Stimuli

Face stimuli were initially selected from a face database created at Bangor University. The Bangor face database comprises photographs of participants with an emotionally neutral expression and self-report measures of various personality and subclinical traits (Kramer and Ward, 2010; Jones et al., 2012; Scott et al., 2013). Individual images were extracted from the database and transformed along four personality or health dimensions (Extraversion, Agreeableness, Neuroticism and Physical Health). These dimensions were chosen because prior work had shown that these dimensions were readily identifiable in composite stimuli, which average faces across multiple identities (Kramer & Ward, 2010).

All face transformations were performed in JPsychomorph (Tiddeman et al., 2001). Face stimuli were produced by transforming an original face image from the database towards an average template of a high trait face (High Trait) or towards an average template of a low trait face (Low Trait). Template faces were produced by creating a composite of the 15 individuals with the highest or lowest ratings along each of the four dimensions. For example, for physical health, an average composite of the 15 most physically healthy individuals in the database was created as well as an average composite of the 15 least physically healthy individuals. This process was repeated for all four dimensions (Extraversion, Agreeableness, Neuroticism and Physical Health). To avoid skin colour or make-up influencing the construction of composite images, only individuals that were white and not wearing make-up were included. Also, to simplify the design space, we only used images of female individuals.

Individual images were selected that were between those included in the high and low composites and also met the above inclusion criteria (i.e., white females who were not wearing make-up). Additionally, the individual in each image had provided consent that their individual face could be shown in later studies. The number of individuals fitting these criteria per trait were: Extraversion = 54, agreeableness = 53, neuroticism = 56, physical health = 54, which made a total of 217 IDs. Note that these were not unique IDs and most were used across traits. An individual face image was then transformed in two ways: towards the high trait composite image by 100% and towards the low trait composite image by 100% (Figure 1). A 100% transform retains the identity cues of the original image whilst shifting the appearance by 100% of the shape, colour, and texture difference between the high and the low composite images. This produced two transformed images per original stimulus (High trait, Low trait), which made 434 images in total.

We transformed stimuli in this manner to exaggerate the distinctive facial features associated with particular trait characteristics, whilst maintaining a variety of facial identities by using individual faces rather than composite images. We did not use composite face images, as this would reduce the variety of identities presented during the scanning task, which may lead participants to disengage. Indeed, we wanted to maintain interest in the stimuli and thus encourage a ‘fresh’ social judgment on every trial and increasing variety of idiosyncratic facial features and identities seemed a concrete way of doing so.

#### Pilot task

To assess the extent to which these stimuli would cue distinct trait judgments, we ran a pilot behavioural experiment (see Supplementary Method). The pilot experiment demonstrated that judgements of Low and High Extraversion, Neuroticism and Physical health were perceived distinctly and as anticipated based on prior research (Kramer & Ward, 2010; Supplementary Figure 2). However, there was no difference in the perception of high and low agreeableness (Supplementary Figure 2). Prior work on agreeableness averaged multiple facial identities to create one composite image (Kramer & Ward, 2010). In the current study, we used individual faces that had been transformed towards High or Low trait features. Therefore, after the pilot study, it was unclear if the lack of distinct behavioural judgments based on agreeableness was due to the method of stimulus construction. We decided to leave the agreeableness stimuli in for the scanning experiment in order to see if the same pattern of results persisted in new participants and, if so, if there were neural effects in the absence of distinct behavioural judgments.

#### Main task

The main task used an event-related design with two types of face stimuli presented (High trait and Low trait faces). The design of the main task is illustrated in Figure 1. Each run comprised 17 blocks of 9 trials. On every trial participants were shown a face and asked to make a social judgement. At the start of each block, participants were shown a written statement and a ratings scale for 4 seconds (1=Strongly disagree, 2= Disagree, 3 = Agree, 4=Strongly agree). The task for participants was to rate how well the person matched the statement. Each trial lasted 3s and participants were instructed to make a judgment based on their initial reaction or “gut instinct”. The scale was always the same, but was included with the statement before each block as a reminder. Participants responded on a button box within the scanner by pressing the corresponding key. Between blocks a white cross was presented on a black screen for a randomly selected duration of 2, 3 or 4 seconds.

Each block contained High and Low versions of stimuli from one category (e.g., Physical Health) and each trial showed a different person. However, participants were not shown high and low versions of the same person in the same category. Instead, participants were shown either a high or a low version of an individual to avoid confusion with seeing the same person transformed to opposite ends of a single dimension. Statements for each block related to the category of stimuli presented in that block. For example, in a physical health block, participants made judgments based on statements concerning physical health. Four statements per category were taken for Extraversion, Agreeableness and Neuroticism from the corresponding scales of the mini-IPIP (Donnellan et al., 2006). An example of an Extraversion statement is “Is the life of the party”. For physical health judgements, items were used from the Short-Form 12-Item Health Survey, which assesses physical health (Ware, Kosinski, & Keller, 1996). An example physical health statement is ‘‘Finds it easy to climb the stairs”. The first block in a run was randomly selected as a starter block. Subsequently, four blocks of each category were presented in a pseudorandom order such that each block followed each other equally often.

Each block began with a starter trial, which was randomly selected from that category. The next 8 trials were sequenced to achieve an even number of novel and repeated trials with novel and repeated trials following each other equally often. Each trial was defined in reference to the preceding trial. For example, a High trait trial that was preceded by a High trait trial would be defined as a repeated trial, whereas a High trait trial that was preceded by a Low trait trial would be defined as a novel trial. This design produced the two conditions of interest, which were modelled as separate regressors in the general linear model: Novel_FaceTrait and Repeated_FaceTrait. The starter trial was included as an additional regressor of no interest since the trial was not preceded by any trial and therefore it was not comparable to the other trials. Each trial was modelled from the onset of the first image for a nominal zero second duration. Across a block there were four trials per condition and across a run there were 68 trials per condition. Each participant completed two runs of the main task, which made 136 trials per condition over the entire experiment. In addition, before entering the scanner, participants completed two practice blocks of the main task.

#### Face localiser

To identify face-selective brain regions, we used an established face localiser (Pitcher et al., 2011). Five categories of stimuli were shown to participants (faces, bodies, scenes, objects, scrambled objects), with one category per block. Each block lasted 18s and showed six 3s movie clips from that category. A total of two blocks were shown in each functional run. At the start, middle and end of each functional run, there was a rest condition for 18s. In the rest condition, a series of six uniform colour fields were presented for 3s each. The order of blocks was reversed from the first to the second bock (e.g., fixation, faces, objects, scenes, bodies, scrambled objects, fixation, scrambled objects, bodies, scenes, objects, faces, fixation). Throughout all blocks, participants were instructed to watch the movies but were not given an explicit task.

#### Theory-of-mind localiser

To localise brain regions associated with ToM, we used an established ToM-localiser (Dodell-Feder et al., 2011; http://saxelab.mit.edu/superloc.php). Participants read 10 short false belief stories, in which the belief characters have about the state of the world is false. Participants also read 10 false photograph stories, where a photograph, map, or sign has out-dated or misleading information. After reading each story, participants had to answer whether the subsequently presented statement is true or false. Each run started with a 12 second rest period, after which the stories (10 seconds) and questions (4 seconds) were presented for 14 seconds combined. Each story was separated by a 12 second rest period. The order of items and conditions was identical for each subject. In the first run, stimuli 1 – 5 from each condition were presented, and the remaining stimuli were presented during the second block.

#### Procedure

Participants completed two runs of the main task. Two additional functional runs were also completed as part of another experiment – one run included a version of an imitation inhibition task (Brass et al., 2000) and one run included a version of a flanker task (Eriksen & Eriksen, 1974). These runs occurred before each run of the main task in order to add variety and offset boredom. Subsequently, participants then completed one run of the face localiser and two runs of the ToM-localiser. The ToM-localiser was always presented after participants had completed the main task, to ensure that participants were not primed towards making trait inferences during the main task. All participants completed an anatomical scan.

### Data acquisition

The experiment was conducted on a 3 Tesla scanner (Philips Achieva), equipped with a 32-channel SENSE-head coil. Stimuli were displayed on a MR safe BOLD screen (Cambridge Research Systems: http://www.crsltd.com/) behind the scanner, which participants viewed via a mirror mounted on the head-coil. T2*-weighted functional images were acquired using a gradient echo echo-planar imaging (EPI) sequence with the following parameters: acquisition time (TR) = 2000 ms; echo time (TE) = 30ms; flip angle = 90°; number of axial slices = 35; slice thickness = 4mm; slice gap = 0.8mm; field of view = 230 × 230 × 167mm^3^. After the functional runs were completed, a high-resolution T1-weighted structural image was acquired for each participant (voxel size = 1 mm^3^, TE = 3.8 ms, flip angle = 8°, FoV = 288 × 232 × 175 mm^3^). Four dummy scans (4 * 2000 ms) were routinely acquired at the start of each functional run and were excluded from analysis. 291 volumes per functional run were collected, except for participant 1 where 288 and 289 volumes were collected in block 1 and 2 respectively.

### Behavioural data analysis

During scanning, faces were rated on four dimensions in a similar manner to the pilot experiment. The four dimensions included Extraversion, Agreeableness, Neuroticism and Physical Health and the ratings scale ranged from 1 to 4 (1 = Strongly disagree, 2 = Disagree, 3 = Agree, 4 = Strongly agree). Ratings on each of these dimensions were compared between high and low transformed stimuli. We expected high transformed stimuli to be rated in a manner that is more consistent with descriptions of the trait category. For instance, based on prior work (Kramer & Ward, 2010), as well as our behavioural pilot data, we would expect stimuli transformed towards high physical health to be rated in a manner consistent with higher physical heath. To compare high and low transformed stimuli, we computed difference scores between high and low stimulus categories as well as interval estimates using 95% confidence intervals (Cumming, 2013). We also computed a paired-samples t-test and a standardised effect size for each difference score (Cohen’s d_z_; Cohen, 1992; Lakens, 2013).

### fMRI data preprocesing and analysis

#### Preprocessing

Head motion was examined for each participant on each task, with an exclusion criteria if displacement across either task exceeded 3 millimetres. We report for each task how many runs or participants were removed for each experiment. fMRI data were analysed with Statistical Parametric Mapping software (SPM8; Wellcome Trust Department of Cognitive Neurology, London, UK: http://www.fil.ion.ucl.ac.uk/spm/). Data were realigned, unwarped, corrected for slice timing, and normalised to the MNI template with a resolution of 3mm^3^. Images were then spatially smoothed (5mm).

#### Analysis

We used spm_ss to perform our primary analyses (Julian et al., 2012; Nieto-Castanon & Fedorenko, 2012; http://www.nitrc.org/projects/spm_ss). Spm_ss enables a subject-specific approach to fMRI data analysis. Like other ROI approaches, functional regions of interest (fROI) are defined and tested in separate data to ensure that the analyses are not circular (Kriegeskorte et al., 2009). The advantage of spm_ss is that it uses an algorithm (or functional parcels from prior datasets) to define fROIs in a group-constrained and subject-specific manner (GSS). This means that the approach benefits from showing group consistency across participants, without requiring complete voxel-level overlap across participants. As such, the approach integrates single-subject specificity within individuals with group-constrained consistency across individuals.

We used GSS to define fROIs using separate localiser data. fROIs were first defined using Face and ToM network localisers before we tested how these fROIs responded in our main task contrasts of interest (RS FaceTraits). To do so, the following steps were taken. 1) Using localiser data, we computed activation maps in individuals, thresholded these images (p < 0.001, uncorrected) and overlaid them on top of one another. The resultant overlay map contains information on the percentage of individuals that show an above threshold response. 2) The overlay map was then divided into regions by an image parcellation algorithm. 3) The resulting regions are then investigated in terms of the proportion of subjects that show some suprathreshold voxels. 4) Regions that overlap in a substantial number of participants (>50%) are then interrogated using independent data (i.e., data from the main task). Statistical tests across participants were performed on percent signal change values extracted from the fROIs according to contrasts of interest.

#### Main task contrasts

For the fMRI data analysis of the main task, we computed our primary contrast of interest: RS Face Traits (Novel_FaceTrait > Repeated_FaceTrait).

#### Face localiser contrasts

Each block was modelled from the onset of the first trial for the entire block (18 seconds). A design matrix was fit for each participant with five regressors per block (Faces, Bodies, Scenes, Objects, Scrambled objects). To identify face-selective regions, a Face > All baseline contrast was evaluated in individual participants (Dynamic Faces > Dynamic Scenes + Objects + Scrambled Objects)

#### ToM localiser contrasts

A design matrix was fit for each participant with 2 regressors, one for each experimental condition (false beliefs and false photographs). The ToM-network was revealed by contrasting false beliefs with false photographs (False Beliefs > False Photographs).

## Results

### Behavioural data

During scanning, high trait faces were rated more consistent with trait characteristics than low trait faces for extraversion t(27)=10.88, p <0.001, d_z_ = 2.06, neuroticism t(27)=4.50, p <0.001, d_z_ = 0.85, and physical health t(27)=3.73, p <0.001, d_z_ = 0.71 (Figure 2). There was no difference between high and low trait faces for judgments of agreeableness t(27)=-0.33, p = 0.63, d_z_ = -0.06 (Figure 2). This pattern of results closely replicates our pilot data.

**Figure 2.**
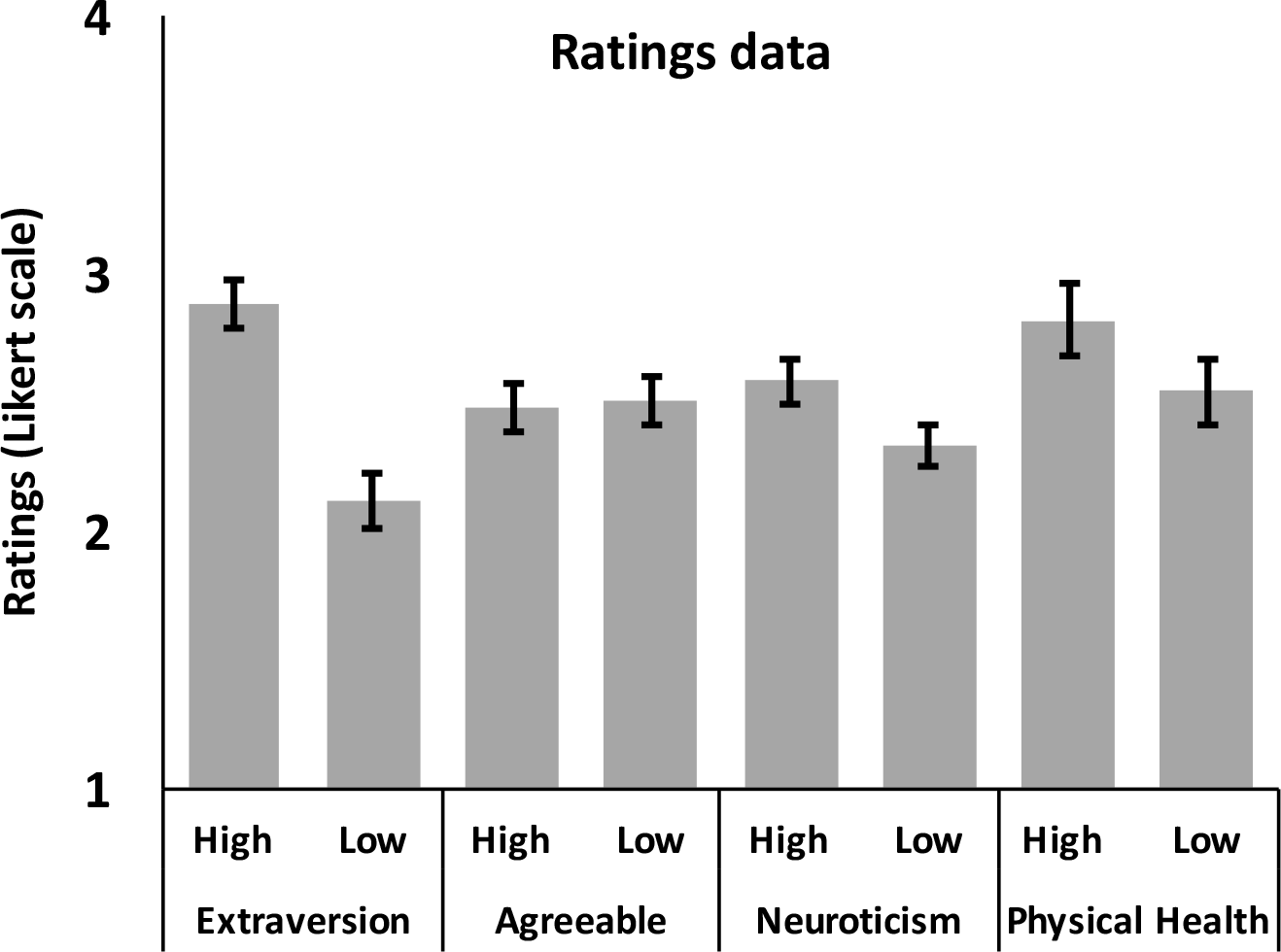
Mean average face ratings during scanning. Error bars are 95% confidence intervals.

### fMRI data

The GSS analysis using the face localiser data revealed nine regions where a majority of participants showed a greater response to faces than all other baseline conditions. Three of these regions are of particular interest given our predictions as they represent the core face perception network. These regions include rOFA, rFFA and r STS/STG. None of the three regions of interest showed RS for Face Traits estimated from data from the main task (Figure 3A; Table 1). If we widen the search to all nine face responsive regions, we do not find RS for Face Traits in any of the ROIs (Supplementary Table 4).

**Figure 3.**
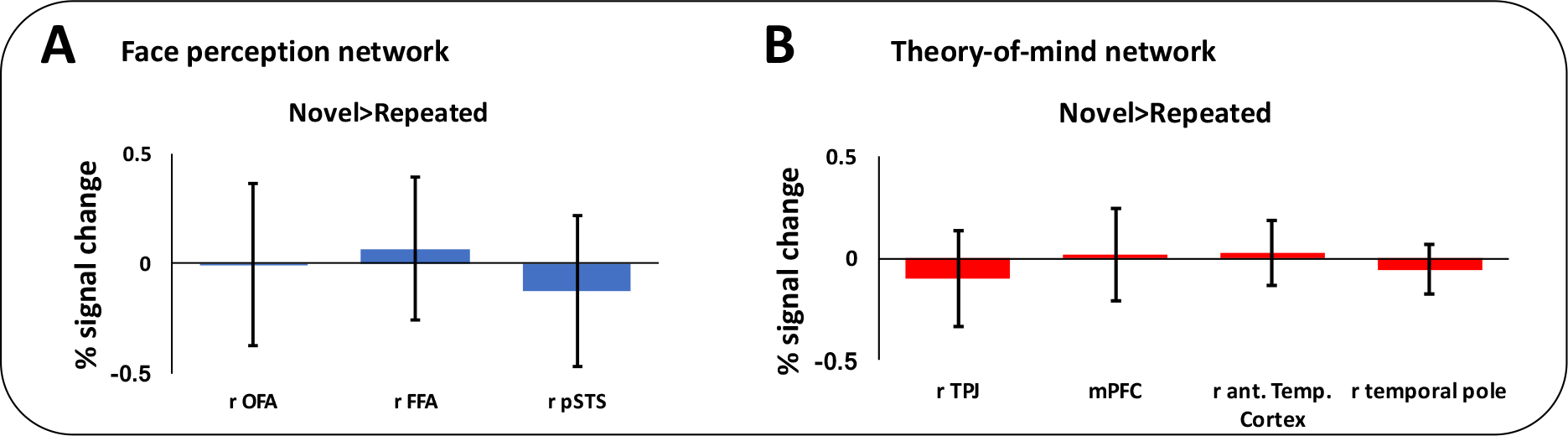
Percent signal change for novel compared to repeated trials in the face perception (A) and theory-of-mind network (B). Error bars are standard error of the mean. Abbreviations: r = right; OFA = occipital face area; FFA = right fusiform face area; pSTS = posterior superior temporal sulcus; TPJ = temporoparietal junction; mPFC = medial prefrontal cortex; ant. Temp. = anterior temporal.

**Table 1.**
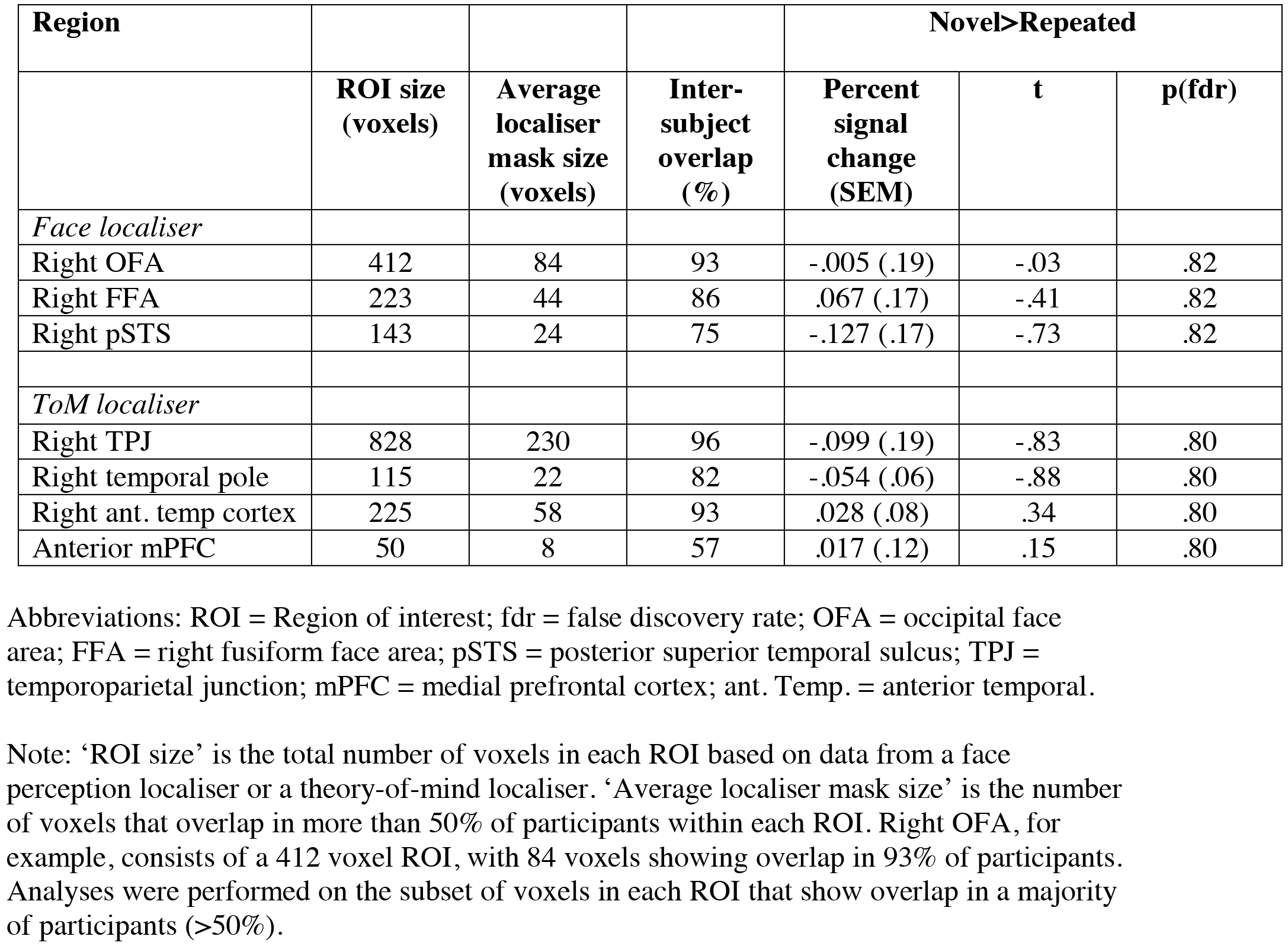
Main task ROI data.

| Region |  |  |  | Novel>Repeated |  |  |
| --- | --- | --- | --- | --- | --- | --- |
|  | ROI size<br>(voxels) | Average<br>localiser<br>mask size<br>(voxels) | Inter-<br>subject<br>overlap<br>(%) | Percent<br>signal<br>change<br>(SEM) | t | p(fdr) |
| *Face localiser* |  |  |  |  |  |  |
| Right OFA | 412 | 84 | 93 | -.005 (.19) | -.03 | .82 |
| Right FFA | 223 | 44 | 86 | .067 (.17) | -.41 | .82 |
| Right pSTS | 143 | 24 | 75 | -.127 (.17) | -.73 | .82 |
| *ToM localiser* |  |  |  |  |  |  |
| Right TPJ | 828 | 230 | 96 | -.099 (.19) | -.83 | .80 |
| Right temporal pole | 115 | 22 | 82 | -.054 (.06) | -.88 | .80 |
| Right ant. temp cortex | 225 | 58 | 93 | .028 (.08) | .34 | .80 |
| Anterior mPFC | 50 | 8 | 57 | .017 (.12) | .15 | .80 |

Abbreviations: ROI = Region of interest; fdr = false discovery rate; OFA = occipital face area; FFA = right fusiform face area; pSTS = posterior superior temporal sulcus; TPJ = temporoparietal junction; mPFC = medial prefrontal cortex; ant. Temp. = anterior temporal.
Note: 'ROI size' is the total number of voxels in each ROI based on data from a face perception localiser or a theory-of-mind localiser. 'Average localiser mask size' is the number of voxels that overlap in more than 50% of participants within each ROI. Right OFA, for example, consists of a 412 voxel ROI, with 84 voxels showing overlap in 93% of participants. Analyses were performed on the subset of voxels in each ROI that show overlap in a majority of participants (>50%).

The GSS analysis using the ToM localiser data revealed nine regions where a majority of participants showed a greater response to false belief stories than false photograph stories. Four of these regions are of particular interest given our predictions regarding specific nodes of the ToM network. These regions include rTPJ, mPFC and r anterior STG / temporal pole. None of these regions showed RS for Face Traits estimated from data from the main task (Figure 3B; Table 1). If we widen the search to all nine regions from the ToM localiser, we do not find RS for Face Traits in any of the ROIs (Supplementary Table 4).

As judgements of agreeableness showed no behavioural differences in perceptions of trait character or health (Figure 2), we removed agreeableness blocks from the analysis, but the results remained the same in both brain networks of interest.

Finally, we completed an exploratory whole-brain analysis, in order to test if regions outside of the Face and ToM networks showed RS for Face Traits. Using SPM8, we calculated Novel > Repeated Face Traits at the single subject level before completing a random effects analysis at the group level using the same contrast. At the group level, no significant clusters of activity were found (p < 0.001, K=10, p<0.05 family wise error corrected). Even at a more liberal threshold (p < 0.001, uncorrected for multiple comparisons), no clusters emerged from this contrast.

Data from this experiment are freely available, including the behavioural and fROI data (osf.io/7knrp), as well as data from the whole-brain analysis (https://neurovault.org/collections/HDLVMPQU/).

## Discussion

Here we show that faces readily cued accurate person judgments regarding extraversion, neuroticism and physical health, but the neural networks associated with face perception and ToM showed no sensitivity in terms of repetition suppression to trait judgements. As such, we do not provide evidence that supports population coding models of face perception that include dimensions for high and low trait features along the ventral visual stream and in the ToM network. Due to aspects of the experimental design and analysis pipeline, which bolstered statistical power and sensitivity, we have reasonable confidence that we could detect effects of a moderate size, should they exist. However, it remains possible that these regions are sensitive to other trait dimensions of person perception such as trustworthiness or other types of facial stimuli, such as synthetic stimuli. The null findings reported here, therefore, add value to models of neural organisation by showing instances where effects are absent or small. In addition, by publishing null results, we provide a less biased scientific record, one that future studies can build upon by appropriately powering studies (Open Science Collaboration, 2015; Simmons et al., 2011). Indeed, future work can use these results to guide further interrogation of what is fundamentally an interesting scientific and social question that relates to understanding the neural mechanisms associated with how trait inferences are cued from facial appearance.

### Understanding the neural basis of impression formation based on facial appearance

The current experiment provides no evidence that populations of neurons in face perception or ToM networks code for facial features that are associated with distinct trait judgements of extraversion, neuroticism or physical health. Moreover, a whole-brain analysis showed no effects in the amygdala or ventral striatum, which have previously been associated with social evaluations of faces based on valence (Bzdok et al., 2011; Mende-Siedlecki et al., 2013; Said et al., 2010; 2011; Todorov et al., 2013). Observers were able to accurately discriminate faces on the basis of the social trait being displayed for the majority of person dimensions. However, we were unable to uncover the neural substrates for this discrimination. In particular, we could not find evidence for our hypothesis that brain regions representing features and judgements for high traits might be separable from those representing low traits. Rather than distinct populations of neurons in the same neural region coding for high and low trait features and judgments, which a neural response consistent with RS would support (Grill-Spector, Henson, & Martin, 2006; Barron et al., 2016), the results may suggest that face perception and ToM networks have a common neural parameter that codes for the perceptual and judgement space under investigation. If so, the same neural populations would be engaged on all trials, whether novel or repeated. For example, if the same features of the face cue high and low judgements, they would be engaged equally on novel and repeated trials. The ultimate judgment would differ between high and low trait faces, but the underlying neural architecture would be similar. This proposal is speculative, however, and would require further testing and confirmation.

An alternative possibility is that RS may not have been sensitive enough to detect the fine-grained population coding structure that was tested. To bolster statistical power, we included a large number of trials per condition for fMRI research (136), we tested 28 participants and we used a single-subject analysis pipeline that has been shown to have relatively high sensitivity and functional resolution in multi-subject analyses (Nieto-Castanon & Fedorenko, 2012). Nonetheless, RS may have been smaller than we could detect with reasonable confidence. Future work may consider multi voxel pattern analysis approaches (Kriegeskorte & Kievit, 2013), which have been shown to be more sensitive than RS approaches in the domain of vision (Sapountzis et al., 2010). In addition, future work may consider the relationship between face and ToM networks as prior functional connectivity research has shown that the ToM network functionally couples with nodes of body perception network (Greven et al., 2016; Greven & Ramsey, 2017a, b). The hypothesis that such future connectivity research could pursue is that the representation of trait judgments from faces may span across face perception and ToM networks rather than only within them.

### Limitations and constraints on generality

In the current study, we do not show RS for trait inferences based on facial appearance. By contrast, other work using written descriptions of behaviour, which imply trait inferences, have shown that vental medial prefrontal cortex (vmPFC) shows RS for trait implying behaviours (Heleven et al., 2017; Heleven & Van Overwalle, 2016; Ma et al., 2014; Van Overwalle et al., 2016). Indeed, this work shows that vmPFC encodes trait representations for familiar (Heleven & Van Overwalle, 2016) and unfamiliar individuals (Heleven et al., 2017), as well as for distinct traits such as valence and competence (Van Overwalle et al., 2016). Therefore, it is important that we acknowledge relevant constraints on the generality of our findings (Simons et al., 2017). Our data, at least with the stimuli that we used, do not support the view that vmPFC stores a person or trait code, which can be easily accessed or engaged irrespective of the type of input (face or text). It could be that written text is simply a more salient way to engage trait inferences, which could lead to the discrepant results. Alternatively, it be might be that not all sources of input (face, text) or all types of person judgment (extraversion, health, trustworthiness) are coded in a similar neural structure. Future work that directly tests interactions between input type and judgments type would be valuable.

Of particular interest for future work would be to test judgments from faces that vary on a valence / trustworthiness dimension (Todorov et al., 2008). In the current study, the behavioural data showed that participants’ judgments did not distinguish between high and low agreeableness faces, which is the closest dimension to valence / trustworthiness. However, participants were sensitive to other dimensions, such as extraversion, neuroticism and physical health. Importantly, recent models of social judgments from faces have shown that appraising faces has three partly distinct dimensions including valence / trustworthiness, dominance and attractiveness (Sutherland et al., 2013). Since judgments of physical health have been associated with attractiveness (Little et al., 2011), our physical health dimension closely resembles a key dimension in the person perception (attractiveness). Therefore, it may be that health and attractiveness judgments, as well as some other types of traits judgment (extraversion, neuroticism), are not coded in the same way as valence / trustworthiness judgments. Indeed, given the role of valence judgments in guiding approach and avoidance behaviours, it may be that there is a more distinct neural architecture dedicated to perceiving such traits.

In the current study, we used morphed images of real human faces. Our findings, therefore, apply most directly to faces that look straightforwardly human. A complementary avenue for future research would be to test models of trait inference from synthetic, computer-generated facial stimuli. The advantage of using computer-generated stimuli would be tighter experimental control, which may boost the ability to detect effects of interest. The obvious disadvantage, however, compared to the current approach of using real photographs, is the artificial limit imposed on ecological validity (Sutherland et al., 2013). Using synthetic images that produce more extreme facial attributes, which differ from the average more, may be important, given research that shows widespread neural responses to faces at high and low ends of continua (Said et al., 2010; 2011). Indeed, even though the majority of trait inferences showed reliable behavioural judgments, it is possible that the similarity between our stimuli reduced the saliency of features that cue trait judgments. Relatedly, we made sure that participants would not see stimuli morphed to different traits in the same block in order to avoid confusion between identities and facial attributes. But, by doing so, this may have made the distinction between high and low exemplars less obvious. An alternative approach would be to show high and low version in the same blocks.

### Open science and the file drawer problem

Since null results and smaller effect sizes are typically relegated to the file drawer (Rosenthal, 1979), the current literature has a publication bias, which prioritises statistically significant results and produces an overestimate of effect sizes. As such, null results from well designed and well powered studies are important if the field is going to move towards a more precise estimate of population effect sizes. Without greater acknowledgement of the value of null results, artificially high estimates of effect sizes will continue to bias models of cognition and brain function, skewing the design of future research and resulting in misallocation of resources (Munafo et al., 2017). Indeed, as outlined above, a null result can make several important contributions to future research (Zwaan et al., 2017). First, replications and extensions can be powered to detect smaller effects or a task can be changed to increase sensitivity. Second, other analysis methods, such as multi-voxel pattern analysis or measures of connectivity (Kriegeskorte & Kievit, 2013; Bullmore & Sporns, 2009), may be prioritised as they may more closely capture the information under investigation. As the data from this study are readily available in online open access repositories, we hope that future research can be guided by this work.

## Acknowledgements

This work was funded by a grant from the Economic and Social Research Council (grant number: ES/K001884/1 to R.R.).

